# Drivers of phyllosphere microbial functional diversity in a neotropical forest

**DOI:** 10.1101/851485

**Authors:** Geneviève Lajoie, Rémi Maglione, Steven W. Kembel

**Affiliations:** Département des sciences biologiques, Université du Québec à Montréal 141, Avenue du Président-Kennedy, Montréal, Qc, Canada, H2X 1Y4

**Keywords:** Microbial communities, Phyllosphere, Functional traits, Host-symbiont matching, Metagenomic shotgun sequencing

## Abstract

**Background:** The phyllosphere is an important microbial habitat but our understanding of how plant hosts drive the composition of their associated leaf microbial communities and whether taxonomic associations between plants and phyllosphere microbes represent adaptive matching remains limited. In this study we quantify bacterial functional diversity in the phyllosphere of 17 tree species in a diverse neotropical forest using metagenomic shotgun sequencing. We ask how hosts drive the functional composition of phyllosphere communities and their turnover across tree species, using host functional traits and phylogeny. We compare functional predictions inferred from 16S gene sequencing with functions estimated from metagenomic shotgun sequencing.

**Results:** Neotropical tree phyllosphere communities are dominated by functions related to the metabolism of carbohydrates, amino acids and energy acquisition, along with environmental signalling pathways involved in membrane transport. While most functional variation was observed within communities, there is non-random assembly of microbial functions across host species possessing different leaf traits. Metabolic functions related to biosynthesis and degradation of secondary compounds, along with signal transduction and cell-cell adhesion were particularly important in driving the match between microbial functions and host traits. These microbial functions were also evolutionarily conserved across the host phylogeny. Functional predictions inferred from 16S gene sequences were weakly correlated with functional annotations from the same samples through metagenomic shotgun sequencing, especially for finer-scale functional annotations.

**Conclusions:** Functional profiling based on metagenomic shotgun sequencing offers evidence for the presence of a core functional microbiome across phyllosphere communities of neotropical trees. While functional turnover across phyllosphere communities is relatively small, the association between microbial functions and leaf trait gradients among host species supports a significant role for plant hosts as selective filters on phyllosphere community assembly. This interpretation is supported by the presence of phylogenetic signal for the microbial traits driving inter-community variation across the host phylogeny. Our comparison of functional annotations derived from 16S genes versus metagenomic shotgun sequencing suggests caution in using functions inferred from 16S genes for studying ecological dynamics in phyllosphere communities. Taken together, our results suggest that there is adaptive matching between phyllosphere microbes and their plant hosts.

## Background

The phyllosphere – the aerial surfaces of plants including leaves – is a widespread microbial habitat that hosts a diversity of microorganisms that play key roles in plant ecology and evolution [1]. Phyllosphere microbes play key roles in plant health [2, 3] and human health [4], and can influence ecosystem function [5]. At a broad taxonomic scale, phyllosphere bacterial communities are consistently dominated by taxa including Actinobacteria, Bacteroidetes, Firmicutes, and Proteobacteria [6], indicating that plants also influence the composition of their microbial partners. A key goal of phyllosphere microbial ecology research has been to identify the adaptive basis of such relationships between plants and associated microbes.

Comparative studies of the taxonomic composition of phyllosphere microbial communities across plant hosts have demonstrated the importance of host identity as a key driver of variation in phyllosphere microbial taxonomic diversity. At fine taxonomic scales, the composition of these communities varies predictably across host plant species [7–9] and across genotypes within host plant species [10, 11]. Plants and associated bacteria also show cophylogenetic associations, with clades of plants and bacteria consistently occurring together [9, 12, 13], suggesting close adaptive associations between plants and their phyllosphere microbes.

Determining whether plant-microbe associations in the phyllosphere have an adaptive basis will require establishing how both plant and microbial functions are related across a range of host species. Plant functional traits – measures of morphology and physiology that capture key axes of variation in plant life history and ecology [14] – have been targeted as a potential proxy for explaining microbial community turnover among plant species. These traits determine the potential for nutrient, metabolite and secondary compound leaching from the plant, which should largely determine the quality of a leaf as a habitat for phyllosphere microbes [15]. In support of this hypothesis, plant functional traits such as leaf mass per area, leaf elemental composition, and growth rate are correlated with phyllosphere microbial community turnover both within [16] and among plant species [12, 17–20].

Several studies have reciprocally identified the broad-scale microbial functional categories and adaptations that epiphytic microbes possess for living on plants [e.g. 16–19]. Functions including the biosynthesis of osmoprotectants such as trehalose and betaine and the production of extracellular polysaccharides are enriched in the phyllosphere and are thought to provide key adaptations to life on leaf surfaces by allowing microbes to attach to the leaf surface and by providing resistance to environmental stresses and plant defenses [25, 26]. However, studies of microbial functions in the phyllosphere have largely been based on comparison of one or a few host plant species. How microbial functions map onto variation in host plant functions in diverse natural communities thus remains largely unknown. As a result, it is not clear whether plant microbiomes exhibit the pattern of taxonomic turnover but functional homogeneity across hosts that has been observed in some animal microbiomes [27] or if a turnover in microbial functions can also be observed across functionally different tree species.

In this study, we quantified the functional repertoire of microbial communities on leaves of multiple tree species in a neotropical forest in Panama using metagenomic shotgun sequencing. We asked which microbial functions are abundant in the phyllosphere, and how these functions are linked to the taxonomy and functional traits of plant hosts. Our central hypothesis was that the plant-microbe taxonomic associations previously observed in this forest [12, 28] should be driven by adaptive matches between microbes and host plants, leading to several key predictions. First, we predicted that microbial functions should vary among host plant species and be correlated with the functional traits of the hosts. Second, we predicted that cophylogenetic associations between trees and microbes should lead to phylogenetic signal in microbial functions present on different plant hosts. Third, we predicted that microbial functions present on leaves should be filtered by the host, since conditions on the leaves of different host plants create a selection pressure on the functions of microbes able to persist on those leaves. Given the increasing interest in using metagenomic predictions of functional genes from metabarcoding data in assessing functional diversity in microbial communities, we lastly aimed to compare the performance of functional predictions from 16S sequencing performed on the same samples in retrieving patterns of functional variation observed in our metagenomic shotgun sequencing dataset [see also 29].

## Results

### Metagenomic shotgun sequencing characterization of phyllosphere microbial functions

Overall, we detected 4587 different functional genes across all samples based on annotation of metagenomic shotgun sequencing of tropical tree phyllosphere communities. Functions related to metabolism were the most abundant overall in our dataset, making up 45% of all functionally annotated sequences (Fig. 1). The principal metabolic functions in the phyllosphere were related to metabolism of amino acids (e.g. amino acid related enzymes), nucleotides (e.g. purine and pyrimidine metabolism), carbohydrates (e.g. pyruvate, glyoxylate and dicarboxylate metabolism), and energy (e.g. oxidative phosphorylation & TCA cycle) (Fig. 1). Groups of functional genes related to environmental and genetic information processing also had a high relative abundance, mainly membrane transport (e.g. transporters), translation (e.g. aminoacyl-tRNA biosynthesis), and signal transduction (e.g. two-component systems).

**Figure 1.**
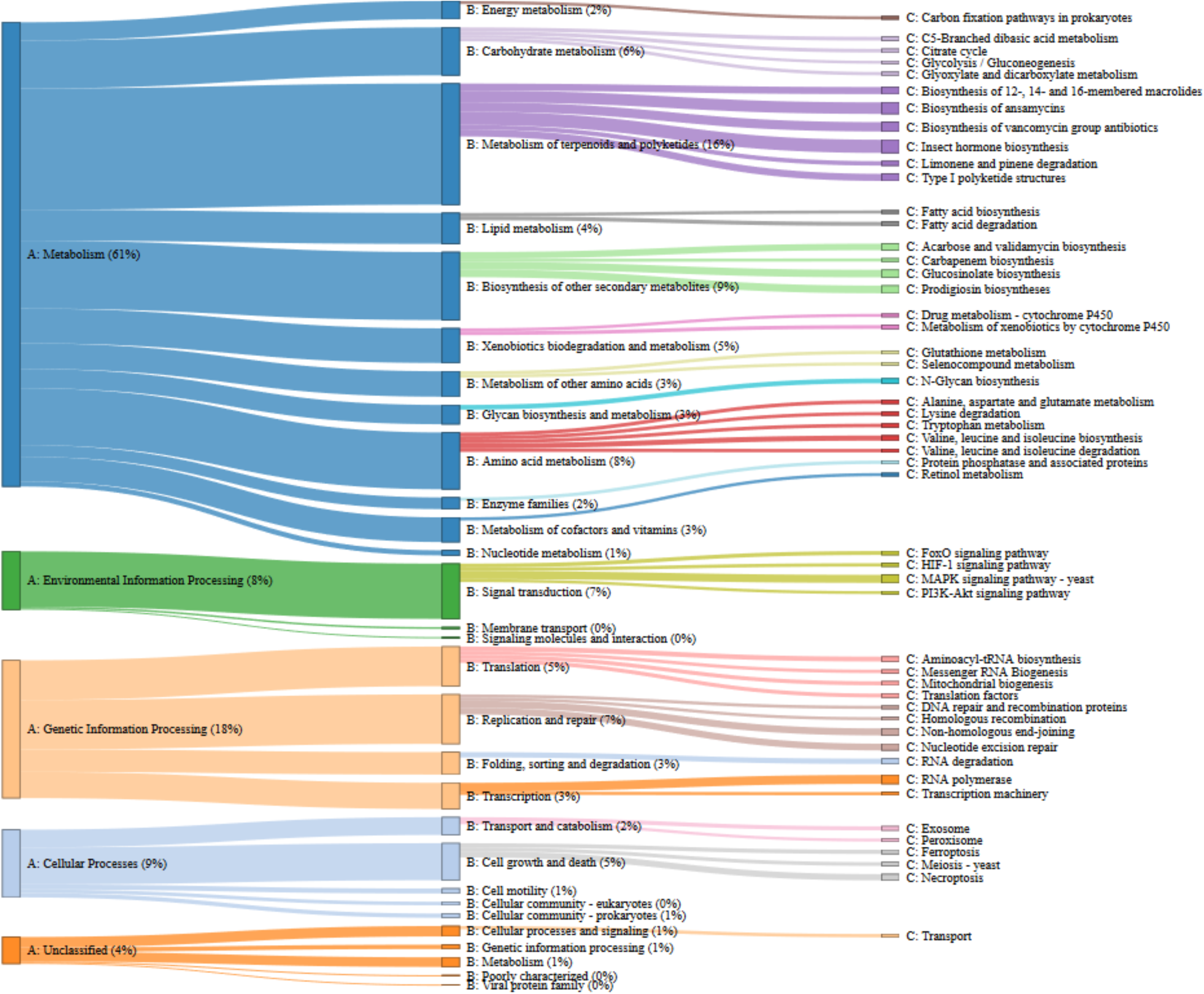
Relative abundance of the most abundant functional pathways detected across 24 tree phyllosphere samples in a neotropical forest in Panama. Functional pathways are classified using the KEGG functional hierarchy [32].

### Variation in phyllosphere functions and taxa among versus within samples

The bacterial functions present on tree leaves were remarkably consistent among different samples. The vast majority of functional variation occurred within samples (>97%), with a very small contribution of functional turnover among samples (<3%) to total functional diversity, regardless of the functional level under study. Most taxonomic diversity was also observed within samples, with a contribution of beta-diversity increasing from 1 to 4.4% of total diversity with a refinement of the taxonomic scale utilized (Table 1). The principal component analysis of bacterial community functional composition indicated that metabolic functions related to biosynthesis and degradation of secondary compounds and antibiotics, as well as functions related to signal transduction and cell-cell adhesion were the most strongly varying among hosts (Fig. 2; Supp. Tab. 1, Additional File 1). We detected 17 Tier 3 functions that exhibited a significantly non-random phylogenetic signal with respect to the host phylogeny (P<0.05; Fig. 3). These functions were mostly involved in the metabolism of terpenoids and polyketides, signal transduction and cellular processes.

**Table 1.** Functional and taxonomic additive diversity partitioning of bacterial communities across 24 tree phyllosphere samples.

|  | Functional |  |  |  |  |  | Taxonomic |  |  |  |  |  |
| --- | --- | --- | --- | --- | --- | --- | --- | --- | --- | --- | --- | --- |
|  | Metagenomic |  |  | 16S |  |  |  |  |  |  |  |  |
|  | Tier 2 | Tier 3 | Functional gene | Tier 2 | Tier 3 | Functional gene | Phylum | Class | Order | Family | Genus | Species |
| Alpha diversity (%) | 100.0 | 99.8 | 97.2 | 99.7 | 99.6 | 99.5 | 99.0 | 99.0 | 99.0 | 98.8 | 98.2 | 95.6 |
| Beta diversity (%) | 0.0 | 0.2 | 2.8 | 0.3 | 0.4 | 0.5 | 1.0 | 1.0 | 1.0 | 1.2 | 1.8 | 4.4 |

**Figure 2.**
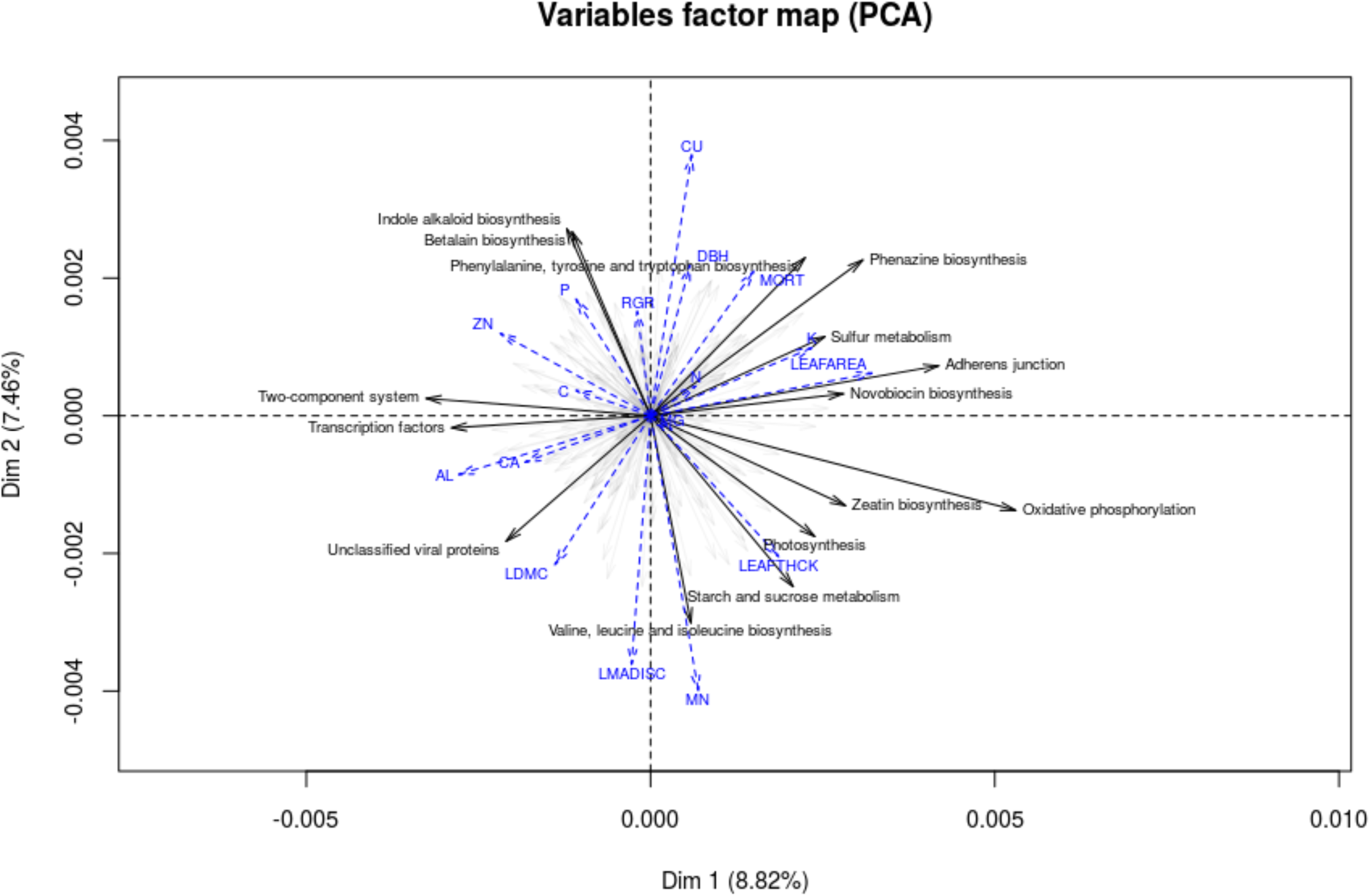
Principal components analysis (PCA) of microbial functional composition from the phyllosphere of neotropical trees. The 20 Tier 3 functions contributing the most to variation among samples are indicated as black arrow. Plant traits were fitted onto the PCA in a configuration that would maximize correlation with the PCA axes and are represented as blue dashed lines. Plant trait abbreviations are the following: Aluminum (AL), Calcium (CA), Carbon (C), Copper (CU), Diameter at breast height (DBH), Leaf area (LEAFAREA), Leaf dry matter content (LDMC), Leaf mass per area (LMA), Leaf thickness (LEAFTHICK), Manganese (MN), Mortality (MORT), Nitrogen (N), Phosphorus (P), Potassium (K), Relative growth rate (RGR), Zinc (ZN).

**Figure 3.**
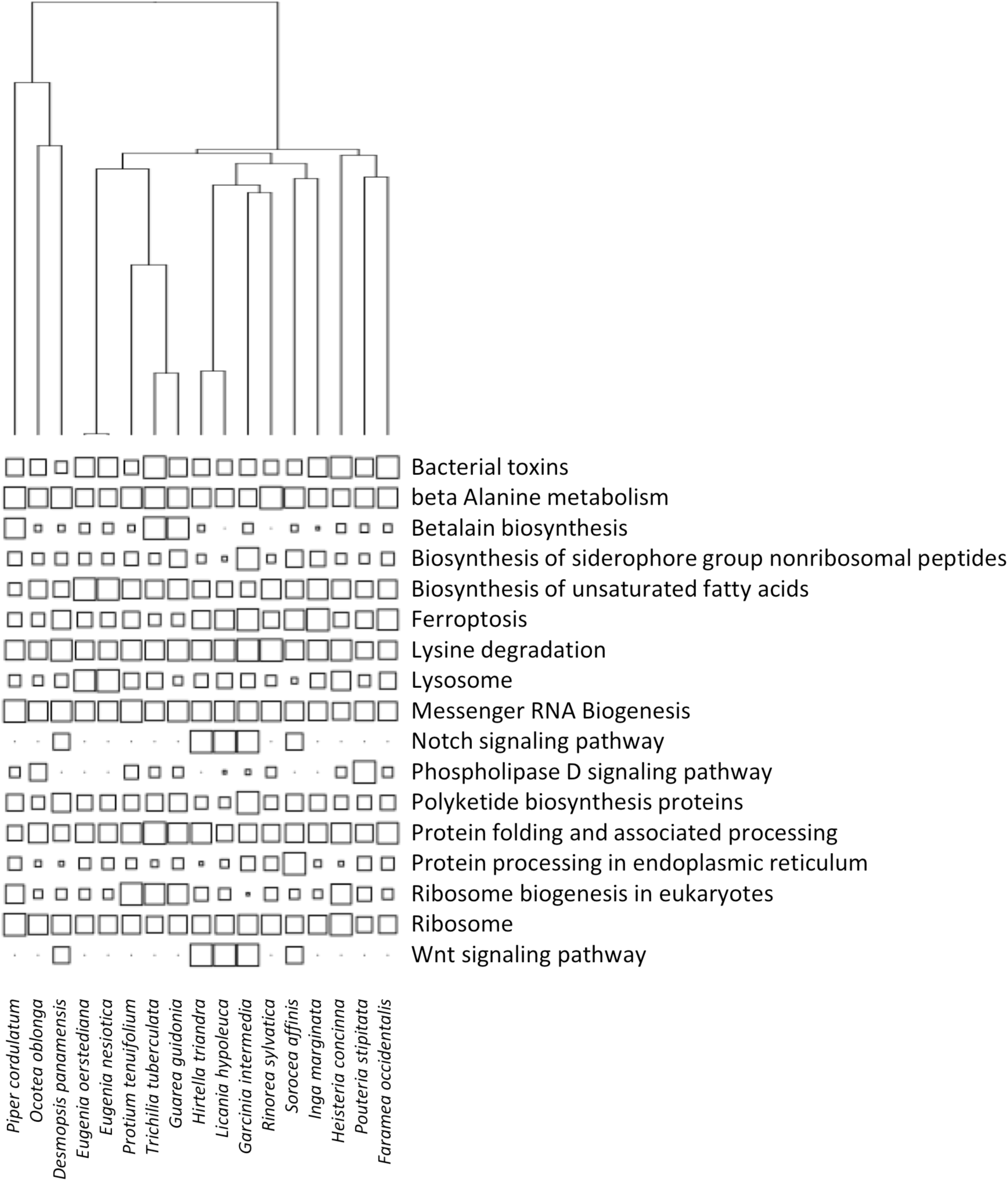
Distribution of microbial functions with respect to plant phylogeny. Distributions are shown for the subset of Tier 3 microbial functions with significant phylogenetic signal according to the K statistic test (P<0.05). Symbol size indicates the scaled relative abundance of microbial functions for each host species.

### Associations between microbial and plant traits and host filtering

Many of the plant traits displayed some level of correlation with the principal axes of microbial functional community composition. Among these, morphological leaf traits (e.g. leaf area, leaf mass per area) were most strongly associated with the first two axes of microbial functional variation. Leaf elemental concentrations of copper, aluminum and manganese were also strongly correlated with these first dimensions. The plant trait gradients explained altogether ∼17% of variation in functional composition among microbial communities. Around half of the microbial Tier 3 functions were significantly more abundant or less abundant than expected by chance in their community, based on a null model keeping both the total abundance of a trait and the number of traits in a community constant (Table 2). The filtering signal was slightly stronger for the microbial taxa than for the microbial functions (Table 2).

**Table 2.**
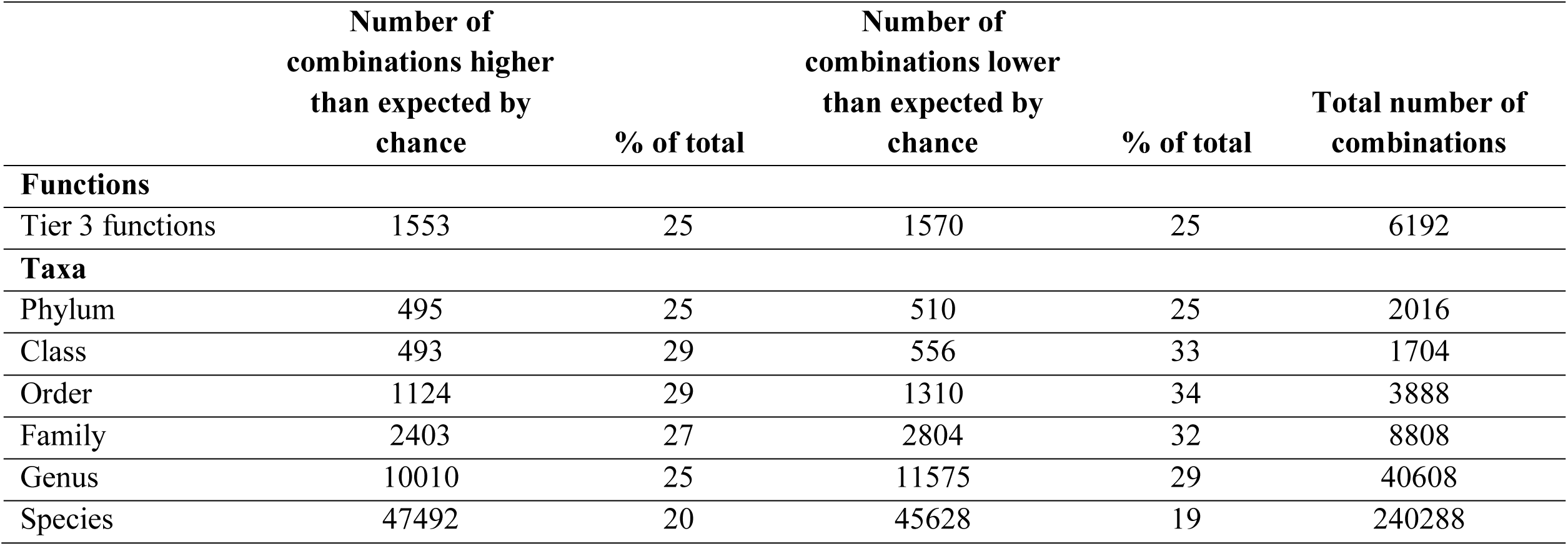
Occurrences of Tier 3 functions and taxa across 24 tree phyllosphere samples.

### Comparison of functional annotations based on metagenomic shotgun versus 16S sequencing

Prediction of the functional content of phyllosphere microbial communities from their 16S rRNA genes using Tax4Fun yielded a higher diversity of functional genes (6429 genes) across all samples compared to predictions from direct annotation of metagenomic shotgun sequence data for the same samples (4587 genes). Most (∼95%) functional genes were detected in both the metagenomic and 16S datasets. The relative abundances of individual genes covaried between the two datasets, with an increasing coherence of functional annotations at broader levels of functional classification (Fig. 4). When testing correlation between metagenomic and 16S annotations within Tier 2 functional categories, we observed generally strong associations (median R^2^ = 0.85) between relative abundances of functions in the two datasets, though the slope of the relationship was often deviant from the 1:1 line (Supp. Fig. 1, Additional File 2). A constrained analysis of principal coordinates analysis revealed that genes related to environmental information processing functions, especially membrane transport and signal transduction, as well as functions related to cellular processes such as bacterial motility proteins and quorum sensing were especially more represented in the 16S dataset, while metabolism functions including nucleotide metabolism and energy metabolism as well as genetic information processing functions (transcription and translation) were more represented in the metagenomic dataset (Supp. Tab. 2, Additional File 1). The type of functional prediction used had an influence on the classification of samples in ordination space. While the classification of samples was consistent among functional levels for each analysis separately, ordinations were incoherent between datasets at all levels (*m^2^* similar to that expected by chance) (Supp. Fig. 2, Additional File 2).

**Figure 4.**
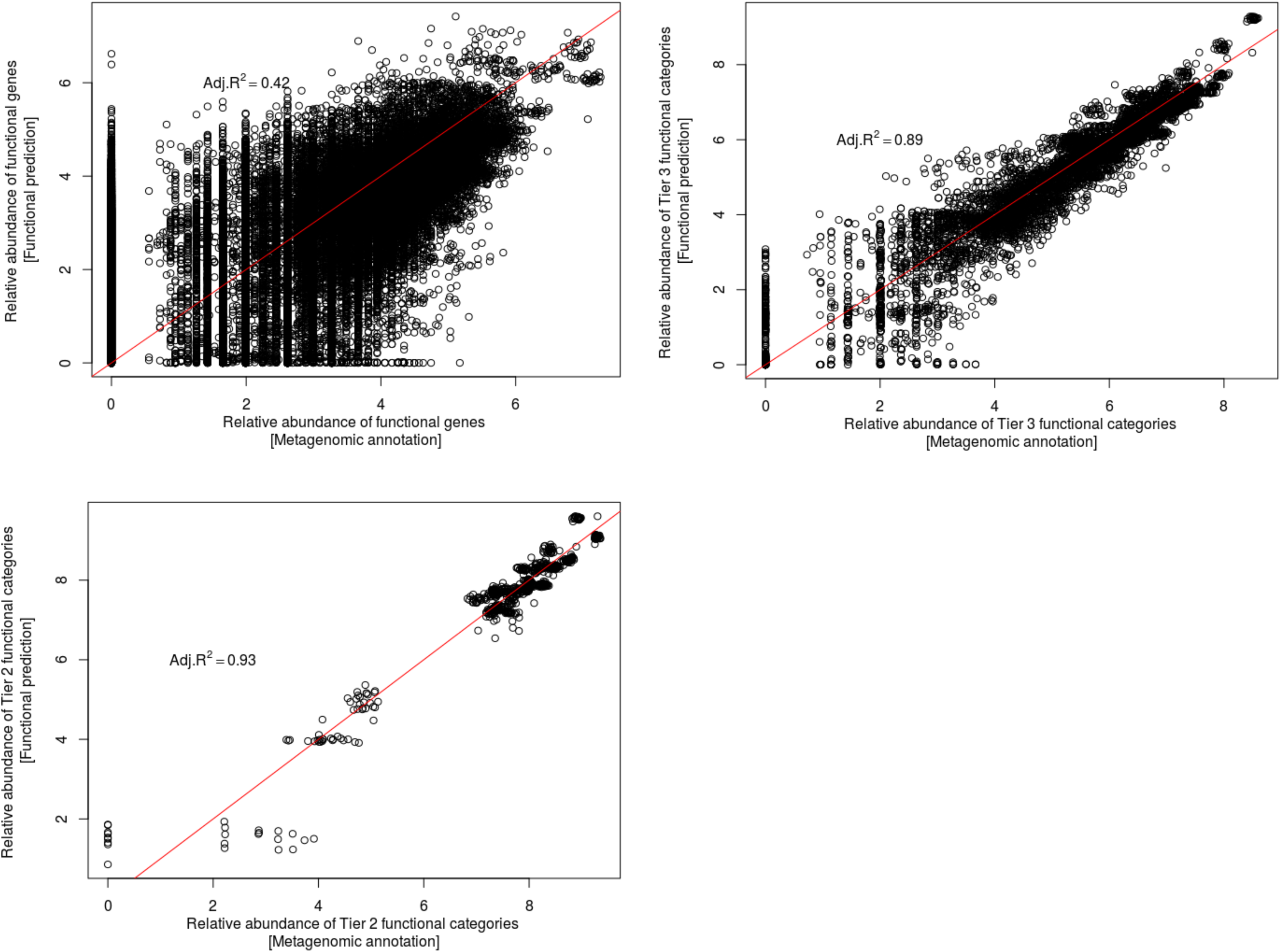
Log-transformed relative abundance of functions detected in the metagenomic annotations and the 16S functional predictions across 24 tree phyllosphere samples in a neotropical forest in Panama. Relative abundances were evaluated at each of 3 functional classification levels. The red line represents a 1:1 relationship between the relative abundances of functions observed at each site, such that points below the red line represent occurrences of functions over-represented in the metagenomic dataset, and those above the red line occurrences of functions over-represented in the 16S dataset.

## Discussion

The functional composition of tree phyllosphere microbial communities in a tropical forest in Panama is largely consistent with those reported in the literature, regardless of the type of plant studied, suggesting the presence of a core functional microbiome in phyllosphere microbial systems. Core functional microbiomes in host-associated systems have also been reported for other hosts. Our study supports findings of an important role for the metabolism of carbohydrates and amino acids in bacterial survival in the phyllosphere [18, 46, 47] that is consistent with the abundance of these compounds in leaf leachates and photosynthates. The main mechanism of energy acquisition from these compounds appeared to be the TCA (citric acid) cycle, as reported in experimental studies of bacterial colonization of the phyllosphere [47]. Membrane transporters were also reported to be an important component of the epiphytic microbe functional repertoire, maximising the ability to monopolize otherwise limiting resources [48]. The abundance of signal transduction functional pathways, involved in the rapid sensing and response to environmental change, would lastly be coherent with the high variability in conditions of humidity, light and temperature in that microbial habitat [25].

The low functional variability in microbiomes observed among tree species represents a further line of evidence supporting the presence of a core phyllosphere functional microbiome. This low variability, observed even at fine functional levels, could be the consequence of essentially similar constraints imposed by the generally harsh leaf environment on its microbial communities, regardless of the specific physiological traits of the host plant species. This low functional turnover among communities was also associated with a low taxonomic turnover, contrasting with reports from phyllosphere-associated temperate systems where species identity was a strong driver of taxonomic composition of the microbial communities [8]. These results could be explained by a finer-scale partitioning of taxa among neotropical than temperate tree species, or a greater overlap in species functional types limiting strong associations between microbial taxa and their hosts. Such differences should be further investigated.

Despite the high levels of convergence in microbial functions among the phyllospheres of different tree species, several lines of evidence support a role for plant species taxonomic and functional identity in driving microbial community assembly. Tree traits explained a notable portion of the functional turnover among microbial communities. Traits correlated with microbial functional turnover (e.g. leaf area, leaf mass per area) are mostly part of the leaf economics spectrum [49], a functional strategy scheme describing photosynthetic resource-use efficiency in plants, which is coherent with what we know of phyllosphere microbial physiology. The ability of a tree to be conservative of its resources and generate thicker and better protected leaves (i.e. high leaf mass per area) is likely to limit the leaching of nutrients from the leaf to the phyllosphere, in turn constituting a filter on resource-use strategies in microbes. The high correlation of leaf mass per area with turnover in microbial communities is coherent with a previously described role for cuticle characteristics in determining functional turnover among leaf microbial communities [16, 20]. The high correlation of aluminum and copper concentrations in leaves with microbial functional variation may be explained by their role as antibiotics. The predominance of two-component systems associated with high aluminum and copper concentrations suggests that the ability to sense and quickly respond to fluxes in these elements at the cell surface might constitute an efficient stress-response to deal with these conditions [50]. This type of plant trait gradient is analog to the leaf chemical gradient described by Yadav and colleagues [51], who reported variation in leaf colonization by phyllosphere microbes on different tree species as a function of their total leaf phenolics content. Taken together, these interpretations are concordant with the importance of energy metabolism, secondary metabolites and antibiotics production as well as environmental sensing in driving functional turnover of microbes among tree species.

Other lines of evidence support the idea that the plant host plays a selective role on microbial community assembly, such as the detection of bacterial traits that are non-randomly structured in the plant phylogeny. While this pattern might arise from the filtering of microbes on phylogenetically structured selective plant traits or from co-evolution of the two partners, it is regardless indicative of an influence of the host on the functional make-up of bacterial phyllosphere communities. Interestingly, the set of pathways that are important in driving functional turnover among communities belong to the same functional categories as the ones that are phylogenetically structured among plant hosts, supporting the proposed match between these bacterial functions and their host’s functional and taxonomic identity. The fact that the relative abundance of a large set of functions was different within communities than that expected by chance given their relative abundance across samples, also supports a role for individual tree species in structuring the functional composition of their phyllosphere bacterial communities. The higher filtering of most microbial taxa relative to microbial functions suggests a role for unmeasured trait variation in driving functional turnover among communities.

The relatively small but significant contribution of functional turnover among microbial communities to the total functional diversity observed across samples suggests that the functions that are of importance in driving the distribution of bacteria across different host trees are actually relatively few compared to those enabling the bacteria to pass the overall “phyllosphere filter” that is needed to survive in the phyllosphere habitat. It remains unknown whether the majority of functional pathways that do not vary among trees are actually important for the ecology of the microbes, or if that trait variation is adaptively neutral within communities. It is also possible that some pathways important for microbial adaptations to leaf physiological gradients are not yet functionally described and are part of the large number of sequences that could not be functionally annotated. Ongoing efforts to better characterize gene functions will help improve the precision of ecological inferences in environmental metagenomes.

The functional predictions generated from annotation of inferred functions from 16S sequences were broadly comparable to those obtained through shotgun metagenomic sequencing at broad functional levels (e.g. Tier 2 functions), but at finer levels (Tier 3 and functional genes) there were numerous discrepancies in the relative abundance of functions inferred using these different approaches. The categories of functions for which we observed the largest discrepancies between the two approaches to functional annotations overlapped with those described for aquatic bacterial communities by Staley and colleagues [29] (i.e. membrane transport, translation). These results point to consistent biases in predictions of metagenomic functions from 16S sequences across ecosystems and warrant caution in interpreting ecological dynamics from inferred functions, especially when Tier 3 or functional gene abundances are being inferred.

## Conclusions

In conclusion, we have identified a core functional microbiome in the phyllosphere of neotropical trees. While most functional variation was observed within individual microbial communities, we reveal a functional matching between the traits of microbes and the traits of plants across 17 tree species, emphasizing the role for energy metabolism, secondary metabolites and antibiotic production as well as environmental sensing in mediating bacterial adaptation to leaf trait gradients in the canopy. Our identification of the adaptive drivers of phyllosphere microbial community composition in this neotropical ecosystem represents a good starting point for identifying the types of microbial traits that could be routinely studied by phyllosphere microbial ecologists to address global questions on the ecological and evolutionary dynamics of phyllosphere microbes. Empirical testing of the fitness consequences of variation in those traits will represent an important next step in understanding adaptive processes in the phyllosphere.

## Methods

### Microbial DNA collection, extraction and sequencing

Microbial communities were collected from the leaves of 24 individual trees from 17 tree species (1-2 samples per species) in the tropical lowland rainforest of Barro Colorado Island, Panama, in December 2010. These samples were selected from a larger pool of samples [12, 28] for which we had sufficient quantities of high-quality DNA, selecting host species to maximize the phylogenetic and functional diversity of hosts. Methodological details of sample collection are described by Kembel et al. 2014 [12]. Briefly, 50-100g of fresh leaves were collected from the subcanopy of one tree of each species. Microbial cells were then washed from each leaf sample using phosphate buffer [1 M Tris•HCl, 0.5 M Na EDTA, and 1.2% CTAB] and collected by centrifuging at 4,000 × g for 20 min. DNA was extracted using MoBio PowerSoil DNA extraction kits and samples stored at −80°C for future analyses. We quantified DNA concentrations and sequenced both extraction negative controls and PCR negative controls for these samples as part of previously published analyses of bacterial 16S and fungal 28S amplicon sequencing of these samples [12, 28]; none of the negative control samples contained measurable concentrations of DNA and upon sequencing they contained fewer DNA sequences than the minimum cut-off for inclusion in analyses As a result, they were all excluded from subsequent analyses in previously published studies and the present study. To quantify the metagenomic structure of each microbial community, we constructed a paired-end metagenomic shotgun library including a random sample of the whole community DNA composition using an Illumina Nextera XT® kit (Illumina reference FC-131-1024). These libraries were then sequenced using Illumina MiSeq paired-end 2 x 250 base pair sequencing (V2 kit, Illumina reference MS-102-2003). Analyses were performed on these 24 samples unless stated otherwise. Results were not influenced by including replicates of the same species (see tests below).

### Microbial taxonomy and functional trait annotation

Metagenomic shotgun sequencing yielded 14,642,408 reads in total. We trimmed sequences to remove Illumina adapters and truncate end-bases with a quality score less than 20, and removed sequences shorter than 25bp, leaving 14,634,072 trimmed and quality-controlled reads. Taxonomic annotation of all sequences in each microbial community was performed to restrict functional analyses only to bacterial sequences. We annotated metagenomic reads using Kaiju, which annotates taxonomic identity of reads by comparing sequenced reads to the microbial subset of the NCBI BLAST non-redundant protein database [30]. Out of the 7,317,036 sequences, we were able to annotate taxonomy to at least the taxonomic level of kingdom for 2,138,885 sequences, of which 2,100,491 sequences were from Bacteria, representing 29% of total sequences. Of these Bacterial sequences, 1,902,749 were annotated to at least the phylum level, representing 26% of total sequences. Analysis of taxonomic composition was carried out on this subset of sequences annotated to at least the bacterial Phylum level. We rarefied all samples to 20,100 randomly chosen sequences per sample for taxonomic composition analyses, resulting in a total of 482,400 sequences for taxonomic analyses.

Functional annotation of microbial sequences was performed via protein homology searches using the KEGG annotation framework [31, 32] via the software COGNIZER [33]. Analyses resulted in the identification of functional genes and categories for 873,082 sequences representing 12% of sequences. In total, of the 7,317,036 bacterial sequences that were obtained from the metagenomic sequencing of all samples, 722,936 sequences were taxonomically annotated as bacteria and had a functional annotation. We lastly classified each of these sequences into functional categories, defined by the BRITE functional hierarchy manually curated for the KEGG annotation system based on published literature [32]. This hierarchy contains four different levels, which were designed as Tier 1, Tier 2, Tier 3 and functional genes, ranging from the more general to the more specific functional assignment (see [29]). Most analyses were performed at the Tier 3 level, in the intent of reaching a balance between the complexity of the data and its interpretability. In a few instances, Tier 3 categories were perfectly correlated across samples so we removed the duplicates from the dataset (Supp. Tab. 3, Additional File 1).

### Inference of microbial functions from 16S gene sequences

Several methods have been proposed to estimate the functional composition of microbial communities based on estimation of functions from 16S gene sequences [e.g. 25–27]. In order to compare estimates of functional composition based on direct sequencing and annotation of metagenomic shotgun sequencing data versus estimation of functions based on 16S sequences, we analyzed previously published 16S gene sequence data for each sample [12]. The randomly rarefied set of 4000 16S sequences per sample described by Kembel et al. [12] were analyzed using Tax4Fun [35] which provided an inferred estimate of the relative abundance of microbial functions in each sample.

### Plant functional traits and phylogeny

We obtained measurements of plant functional traits for all plant species from a dataset collected previously on Barro Colorado Island [37]. This trait database initially included 21 whole-plant and leaf traits, but we reduced these traits to a subset of 12 traits with limited overlap in functional significance [38]. This reduced set of traits included height at maturity, sapling growth rate and sapling mortality rate as whole-plant resource-use traits, leaf area and leaf dry matter content as leaf structural traits, and a suite of leaf elemental chemistry traits including concentration of aluminum, calcium, copper, magnesium, phosphorus, zinc and nitrogen content. A phylogenetic hypothesis for host plant species was obtained by grafting tree species onto a dated megatree of angiosperms provided by Zanne et al. [39] using Phylomatic v.3 [40].

### Variation in phyllosphere functions among versus within samples

We determined the contributions of within- and among-sample variation in function of total functional variation among metagenomic samples using additive diversity partitioning implemented in the R package entropart [41]. The proportions of alpha and beta diversity were calculated as a ration of the portion of alpha (or beta) diversity on total diversity. Analyses were performed at three levels of functional aggregation (Tier 1 to Tier 3). We tested whether the presence of two samples rather than one for some of the sampled species would affect this diversity partitioning by subsampling the dataset to include all possible combinations of samples totally a single sample per species (n=128) and rerunning the analyses. This subsampling did not affect our results (Supp. Fig. 3, Additional File 2), such that we kept the 24 samples in the subsequent analyses. We then compared sources of turnover for functions and taxonomy between samples by performing the same analysis from the taxonomically annotated metagenomic sequences, defined at levels from phylum to species.

### Associations between microbial and plant traits

We performed a principal component analysis (PCA) of functional trait matrices and identified the functions contributing most to variation along the first axes of variation using R package FactoMineR [42]. We fitted the plant traits onto this ordination to identify correlations between bacterial traits driving the PCA and the plant traits. We evaluated the influence of tree species replicates in our samples on these results and did not uncover important differences in the main drivers of functional differences among samples when excluding these duplicates such that all 24 samples were kept in this analysis.

We quantified the phylogenetic signal in associations between microbial functions and host plant phylogeny using function *multiphylosignal* from R package Picante [43] to calculate Blomberg’s K and an associated P-value, which quantifies whether a microbial function exhibits stronger phylogenetic signal than expected by chance. We selected a single random sample per host species for those host species with more than one sample prior to calculating phylogenetic signal. We repeated this for different random subsamples and it did not qualitatively change the results so we report phylogenetic signal for a representative random subsample.

### Host filtering of microbial functions and taxa

The degree of host filtering on microbial communities was assessed by comparing the occurrence of traits in observed communities to those obtained from 999 randomizations of community trait matrices. Host filtering was detected as an over- or under-representation of the given trait in individual communities. Randomizations were generated by permutations of the trait matrix preserving row and column totals. For each site and bacterial trait combination, we compare the observed frequency of the trait to the random values to assess whether it was lower or higher than expected by chance. To compare the strength of functional vs. taxonomic filtering, we applied the same procedure to the taxonomic datasets defined at each of six taxonomic levels, from the phylum to the species.

### Comparison of functional annotations based on shotgun metagenomic versus 16S sequencing

We compared functional annotations obtained through shotgun metagenomic sequencing with those obtained from functional predictions made from 16S sequencing on the same samples. Since one sample was an outlier in the 16S functional predictions so we removed it in both datasets prior to analyses, resulting in 23 samples total.

We first compared the relative abundances of individual functional pathways across the two datasets by performing a correlation analysis between their average abundance across all samples. The functional pathways the furthest from the 1:1 relationship would be the most over- or under- represented in either method. We then determined the functional pathways which were the most important in differentiating between the metagenomic and the 16S functional annotations by performing a Constrained Analysis of Principal Coordinates (CAP) analysis on Bray-Curtis distances. We next tested whether such differences would have an importance in the ecological classification of samples by performing a Procrustes analysis on the functional tables of each dataset [44], and tested the degree of similarity in the relationships among sites calculated for each dataset using a permutation approach. The degree of similarity is described by the *m^2^* term, representing the sum of the squared deviations between sample positions in one dataset vs. the other. A *m^2^*statistic smaller than expected by chance indicates that the two datasets are more similar than expected by chance [45]. Finally, we generated a diversity partitioning of the 16S functional predictions as described above for the metagenomic functional annotations to determine the impact of the 16S prediction approach in the description of biodiversity within and across samples.

## Declarations

### Ethics approval and consent to participate

Not applicable.

### Consent for publication

Not applicable.

### Availability of data and material

The datasets generated and/or analysed during the current study are available in a MG-RAST repository: https://www.mg-rast.org/linkin.cgi?project=mgp91848

The scripts used to perform analyses for the current study are available in a GitHub repository: https://github.com/glajoie1/panama_metagenome

### Authors’ contributions

GL and SK conceptualized the study. SK collected the data. RM and SK curated the data. GL and SK analyzed the data and wrote the manuscript. All authors revised and accepted the final manuscript.

## Acknowledgements

We thank Travis Dawson and Luis Tovar for assistance with sample processing and sequencing.

**Additional Files Additional File 1 (.docx)**

Supplementary Tables. This additional file contains 3 supplementary tables, referred to in the main text.

**Additional File 2 (.docx)**

Supplementary Figures. This additional file contains 3 supplementary figures, referred to in the main text.

**Additional File 3 (.newick)**

Host phylogeny. A phylogenetic hypothesis for host plant species obtained by grafting tree species from the study site onto a dated megatree of angiosperms (see Methods for details).

