## Additional File 1 for "Drivers of phyllosphere microbial functional diversity in a neotropical forest"

**Supplementary Tables**

**Supp. Tab. 1.** The 25 Tier 3 functions contributing the most to variation among samples. Columns correspond to functional categories defined by the Kegg hierarchy (see Methods).

| **Tier 1**  **Functional category** | **Tier 2**  **Functional category** | **Tier 3**  **Functional category** |
| --- | --- | --- |
| Cellular Processes | Cell growth and death | Apoptosis |
| Cellular Processes | Cellular community - eukaryotes | Adherens junction |
| Cellular Processes | Transport and catabolism | Exosome |
| Environmental Information Processing | Signal transduction | cGMP - PKG signaling pathway |
| Environmental Information Processing | Signal transduction | MAPK signaling pathway - fly |
| Environmental Information Processing | Signal transduction | Two-component system |
| Genetic Information Processing | Replication and repair | DNA repair and recombination proteins |
| Genetic Information Processing | Transcription | Transcription factors |
| Metabolism | Amino acid metabolism | Phenylalanine, tyrosine and tryptophan biosynthesis |
| Metabolism | Amino acid metabolism | Valine, leucine and isoleucine biosynthesis |
| Metabolism | Biosynthesis of other secondary metabolites | Betalain biosynthesis |
| Metabolism | Biosynthesis of other secondary metabolites | Indole alkaloid biosynthesis |
| Metabolism | Biosynthesis of other secondary metabolites | Isoquinoline alkaloid biosynthesis |
| Metabolism | Biosynthesis of other secondary metabolites | Novobiocin biosynthesis |
| Metabolism | Biosynthesis of other secondary metabolites | Phenazine biosynthesis |
| Metabolism | Biosynthesis of other secondary metabolites | Phenylpropanoid biosynthesis |
| Metabolism | Biosynthesis of other secondary metabolites | Tropane, piperidine and pyridine alkaloid biosynthesis |
| Metabolism | Carbohydrate metabolism | C5-Branched dibasic acid metabolism |
| Metabolism | Carbohydrate metabolism | Citrate cycle (TCA cycle) |
| Metabolism | Carbohydrate metabolism | Starch and sucrose metabolism |
| Metabolism | Energy metabolism | Oxidative phosphorylation |
| Metabolism | Energy metabolism | Photosynthesis |
| Metabolism | Energy metabolism | Sulfur metabolism |
| Metabolism | Metabolism of cofactors and vitamins | Pantothenate and CoA biosynthesis |
| Metabolism | Metabolism of terpenoids and polyketides | Zeatin biosynthesis |
| Unclassified | Genetic information processing | Replication, recombination and repair proteins |
| Unclassified | Viral protein family | Unclassified viral proteins |

**Supp. Tab. 2.** Tier 3 functions most strongly associated with the first axis of a Constrained Analysis of Principal Coordinates, emphasizing the differences between the functional composition of communities defined using the 16S functional predictions and the metagenomic functional annotations.

| **Tier 1**  **Functional category** | **Tier 2**  **Functional category** | **Tier 3**  **Functional category** | **Correlation coefficient** | **Overrepresented in dataset** |
| --- | --- | --- | --- | --- |
| Environmental Information Processing | Membrane transport | Transporters | -0.487 | 16S functional predictions |
| Environmental Information Processing | Signal transduction | Two-component system | -0.253 | 16S functional predictions |
| Metabolism | Metabolism of terpenoids and polyketides | Polyketide biosynthesis proteins | -0.191 | 16S functional predictions |
| Cellular Processes | Cell motility | Bacterial motility proteins | -0.166 | 16S functional predictions |
| Cellular Processes | Cellular community - prokaryotes | Quorum sensing | -0.147 | 16S functional predictions |
| Metabolism | Metabolism of terpenoids and polyketides | Nonribosomal peptide structures | -0.138 | 16S functional predictions |
| Genetic Information Processing | Transcription | Transcription factors | -0.131 | 16S functional predictions |
| Metabolism | Metabolism of terpenoids and polyketides | Biosynthesis of siderophore group nonribosomal peptides | -0.125 | 16S functional predictions |
| Metabolism | Glycan biosynthesis and metabolism | Glycosaminoglycan binding proteins | -0.124 | 16S functional predictions |
| Metabolism | Enzyme families | Protein kinases | -0.116 | 16S functional predictions |
| Genetic Information Processing | Transcription | RNA polymerase | 0.096 | Metagenomic annotations |
| Genetic Information Processing | Replication and repair | DNA repair and recombination proteins | 0.097 | Metagenomic annotations |
| Genetic Information Processing | Translation | Aminoacyl-tRNA biosynthesis | 0.103 | Metagenomic annotations |
| Genetic Information Processing | Translation | Mitochondrial biogenesis | 0.104 | Metagenomic annotations |
| Metabolism | Nucleotide metabolism | Pyrimidine metabolism | 0.122 | Metagenomic annotations |
| Metabolism | Energy metabolism | Oxidative phosphorylation | 0.132 | Metagenomic annotations |
| Genetic Information Processing | Translation | Ribosome | 0.134 | Metagenomic annotations |
| Metabolism | Nucleotide metabolism | Purine metabolism | 0.141 | Metagenomic annotations |
| Unclassified | Poorly characterized | Function unknown | 0.161 | Metagenomic annotations |
| Cellular Processes | Transport and catabolism | Exosome | 0.163 | Metagenomic annotations |

**Supp. Tab. 3.** Tier 3 functional categories that are perfectly correlated across samples. The category on the left-hand column was kept for all analyses, while the correlated categories in the right-hand column were discarded.

| **Functional category in dataset** | **Correlated functional category removed from dataset** |
| --- | --- |
| Adherens junction | Focal adhesion  Hippo signaling pathway -fly  Hippo signaling pathway  Phagosome  Rap1 signaling pathway  Regulation of actin cytoskeleton  Tight junction |
| Biosynthesis of 12-, 14- and 16-membered macrolides | Type I polyketide structures |
| Biosynthesis of type II polyketide backbone | Tetracycline biosynthesis |
| Endocytosis | Ras signaling pathway |
| Glycosphingolipid biosynthesis - ganglio series | Various types of N-glycan biosynthesis |
| NF-kappa B signaling pathway | TNF signaling pathway  VEGF signaling pathway |
| Notch signaling pathway | Wnt signaling pathway |
