## Additional File 2 for "Drivers of phyllosphere microbial functional diversity in a neotropical forest"

**Supplementary Figure Legends**

**Supp. Fig. 1.** Relative abundance of functions detected in the metagenomic annotations (x axis) and the 16S functional predictions (y axis), within Tier 2 functional categories (A-E). The red line represents a 1:1 relationship between the relative abundances of functions observed at each site, such that points below the red line represent occurrences of functions over-represented in the metagenomic dataset, and those above the red line occurrences of functions over-represented in the 16S dataset.

**Supp. Fig. 2.** Procrustes analyses of 16S functional predictions and metagenomics functional annotations, for three different levels of functional categorizations. In all cases, the *m^2^* metric was no smaller than expected by chance (999 permutations).

**Supp. Fig. 3.** Distribution of alpha, beta and gamma diversities generated from 128 subsampling of the metagenomic functional dataset to include only one sample per tree species. Despite variation observed among bootstraps, the relative importance of alpha vs. gamma diversity stayed constant at 97.3% alpha diversity and 2.7% beta-diversity for all subsamples. The red vertical line indicates the observed value.

A) Genetic Processes

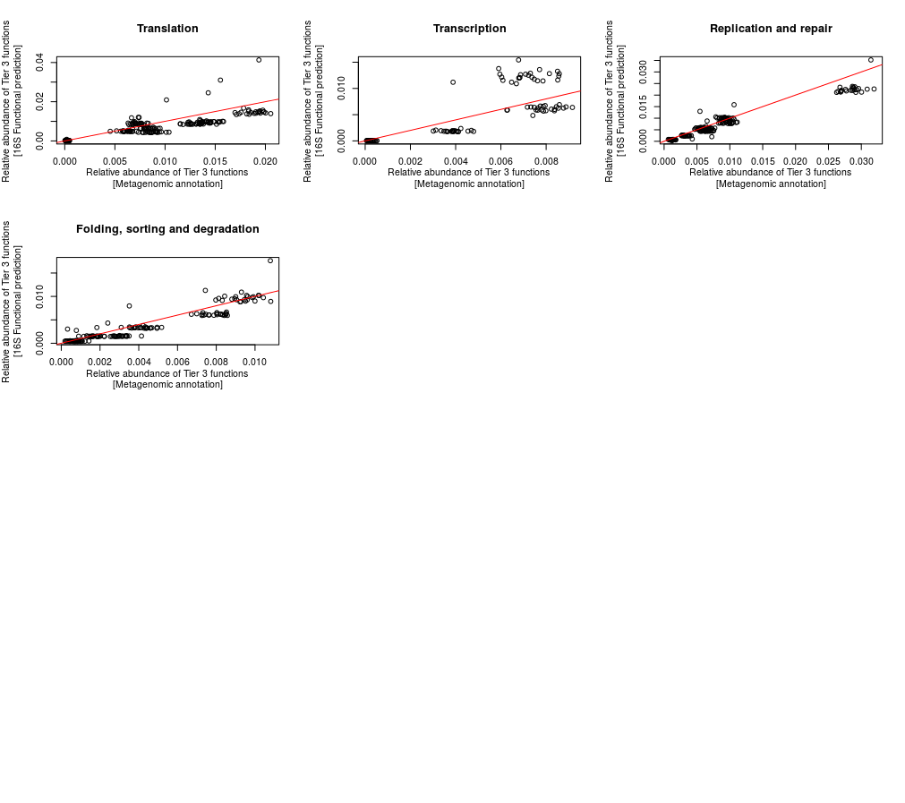

B) Environmental signaling

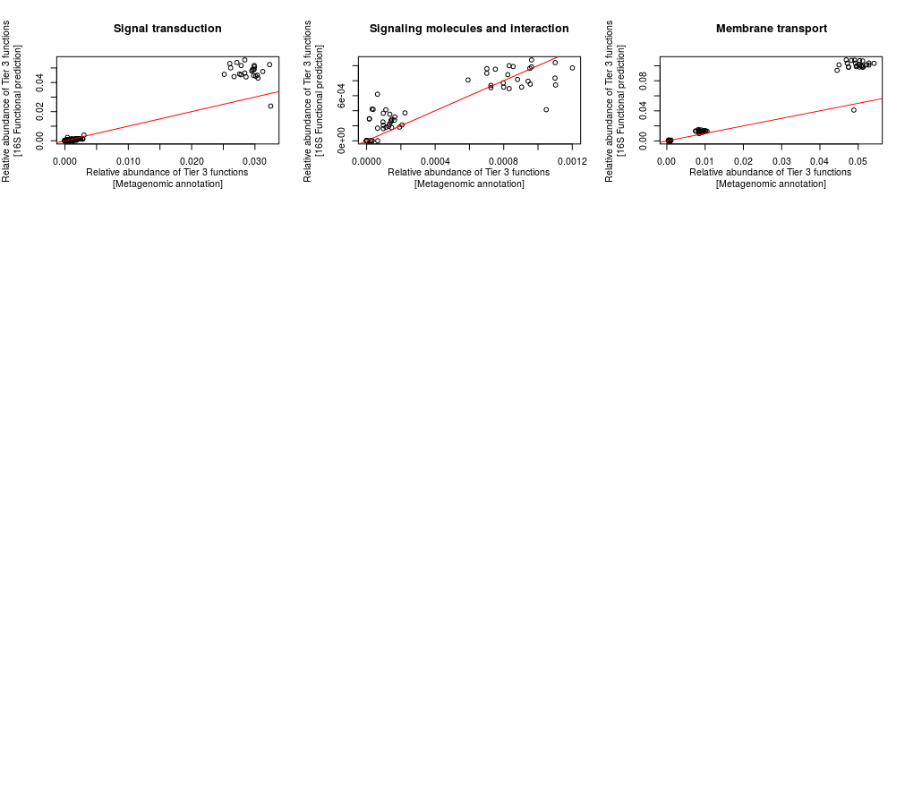

**Supp. Fig. 2 (A-B)**

C) Unknown

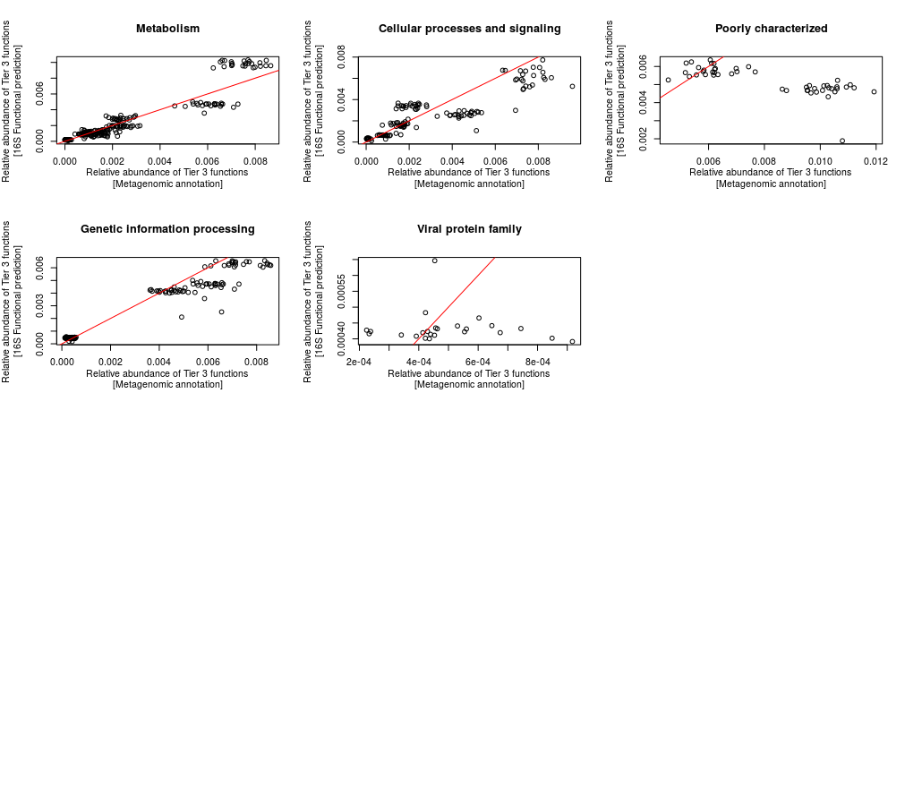

**Supp. Fig. 2 (C)**

D) Cellular processes

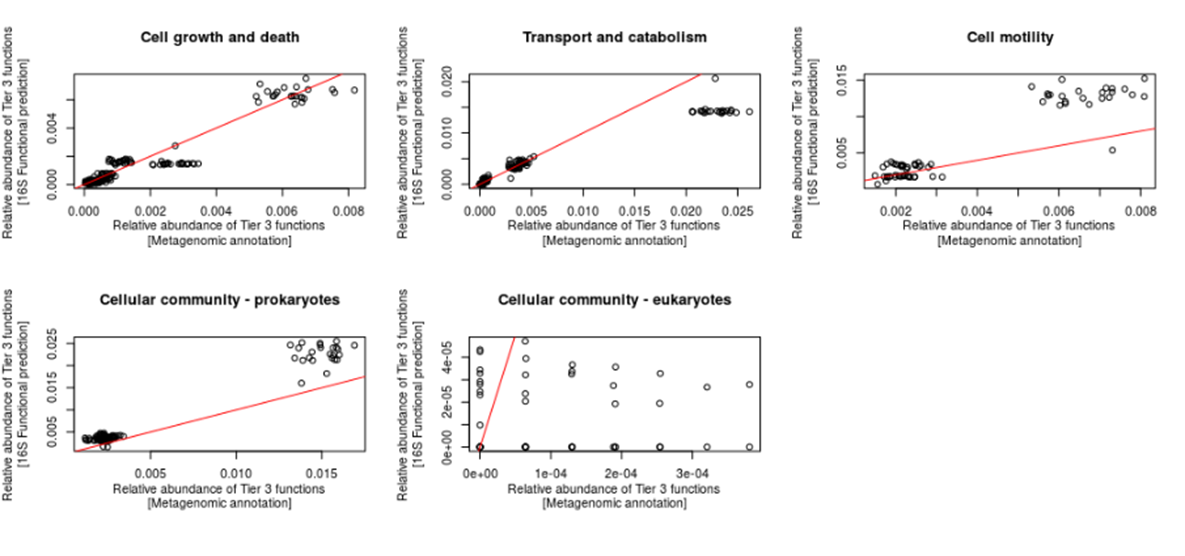

**Supp. Fig. 2 (D)**

E) Metabolism

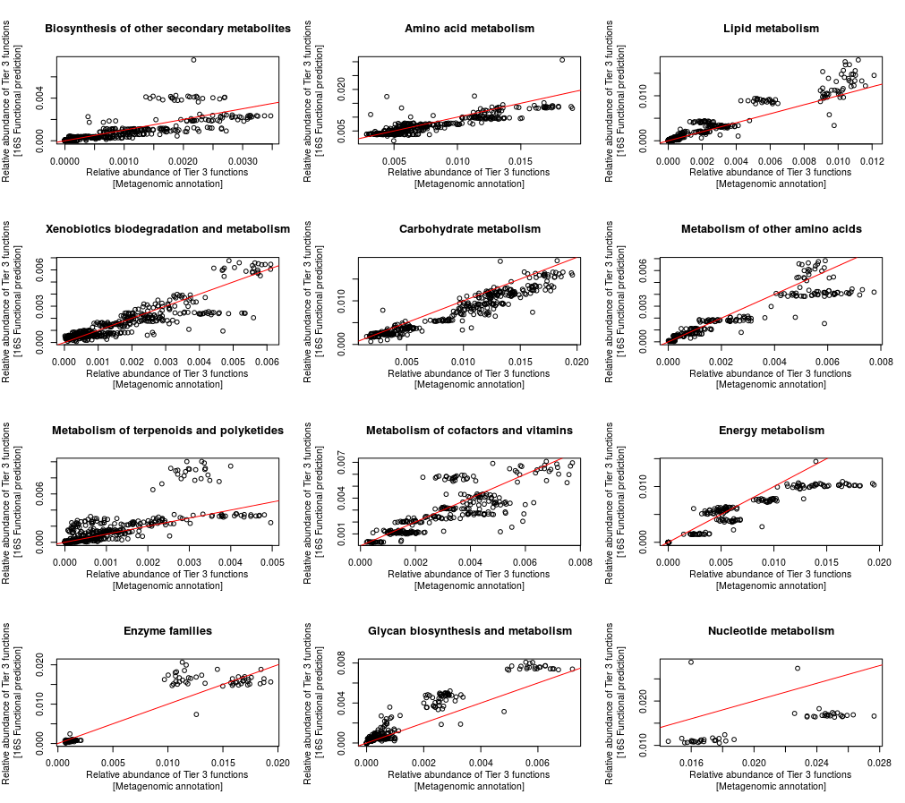

**Supp. Fig. 1 (E)**

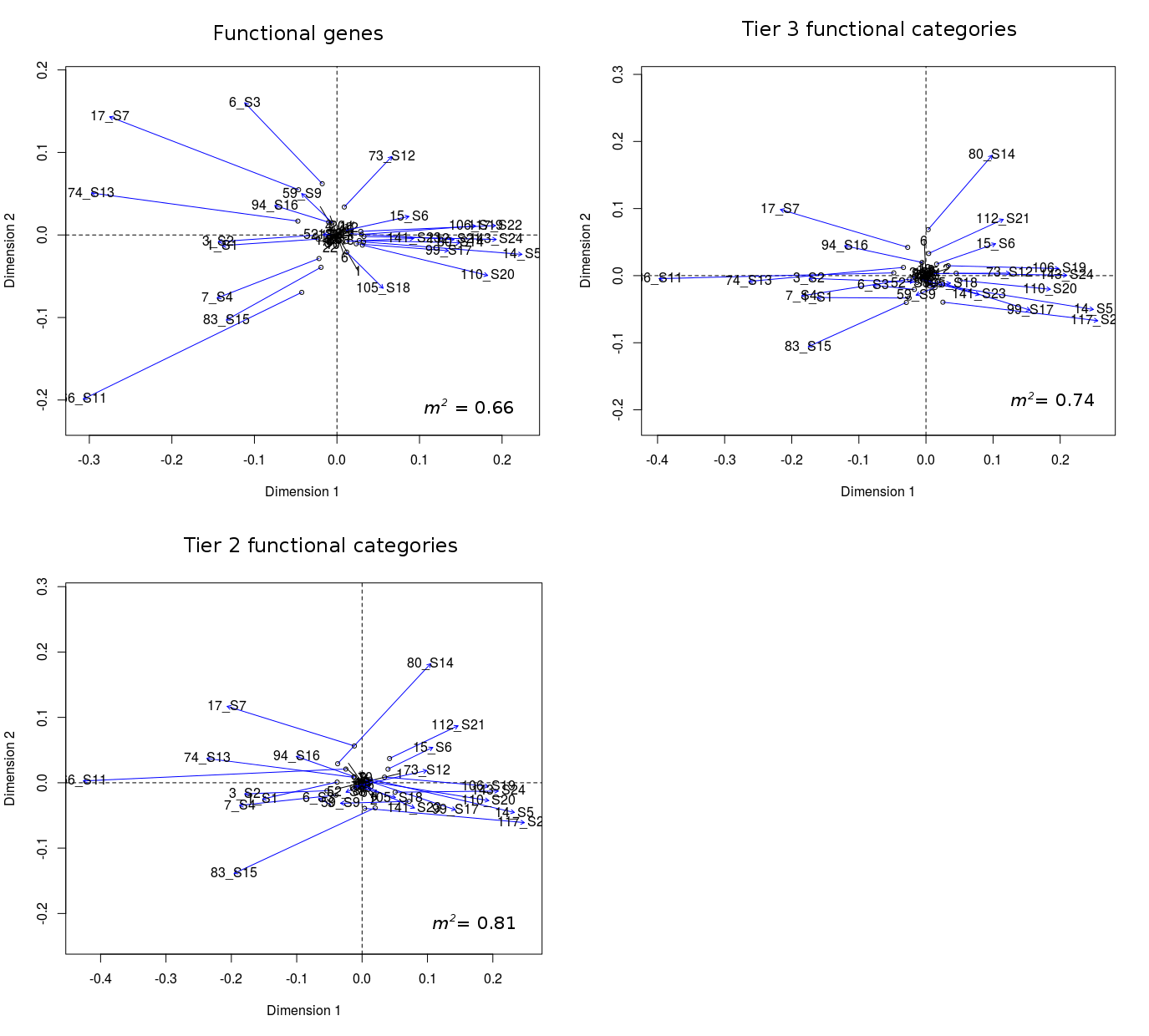

**Supp. Fig. 2.**

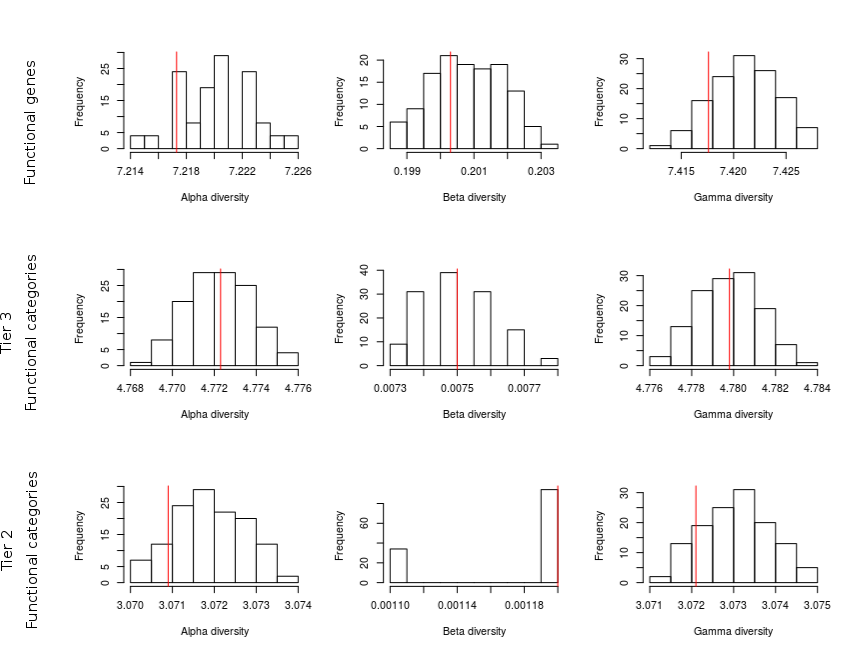

**Supp. Fig. 3.**
